# Using a biologically mimicking climbing robot to explore the performance landscape of climbing in lizards

**DOI:** 10.1101/2021.01.12.426469

**Authors:** Johanna T. Schultz, Hendrik K. Beck, Tina Haagensen, Tasmin Proost, Christofer J. Clemente

**Author notes:** **Author Contributions** JTS contributed towards robot design, data collection, analysis, and co-wrote the manuscript. HKB contributed towards robot design, data collection, and analysis. TH and TP contributed toward data collection. CJC conceived the idea, contributed to robot design, data collection, analysis, and co-wrote the manuscript.

## Abstract

The life and death of an organism often depends on its ability to perform well at some ecologically relevant task. Yet despite this significance we have little idea how well species are optimised for competing locomotor tasks. Most scientists generally accept that the ability for natural systems to become optimised for a specific task is limited by structural, historic or functional constraints. Climbing lizards provide a good example of constraint where climbing ability requires the optimization of conflicting tasks such as speed, stability, or efficiency. Here we reconstruct multiple performance landscapes of climbing locomotion using a 10-DOF robot based upon the lizard bauplan, including an actuated spine, shoulders, and feet, the latter which interlock with the surface via claws. This design allows us to independently vary speed, foot angles, and range of motion, while simultaneously collecting data on climbed distance, stability and efficiency. We first demonstrate a trade-off between speed and stability with high speeds resulting in decreased stability and low speeds an increased cost of transport. By varying foot orientation of fore and hindfeet independently, we found geckos converge on a narrow optimum for both speed and stability, but avoid a secondary wider optimum highlighting a possible constraint. Modifying the spine and limb range of movement revealed a gradient in performance. Evolutionary modifications in movement among extant species appear to follow this gradient towards areas which promote speed and efficiency. This approach can give us a better understanding about locomotor optimization, and provide inspiration for industrial and search-and-rescue robots.

**Significance Statement:** Climbing requires the optimization of conflicting tasks such as speed, stability, or efficiency, but understanding the relative importance of these competing performance traits is difficult.

We used a highly modular bio-inspired climbing robot to reconstruct performance landscapes for climbing lizards. We then compared the performance of extant species onto these and show strong congruence with lizard phenotypes and robotic optima.

Using this method we can show why certain phenotypes are not present among extant species, illustrating why these would be potentially mal-adaptive.

These principles may be useful to compare with relative rates of evolution along differing evolutionary histories. It also highlights the importance of biological inspiration towards the optimization of industrial climbing robots, which like lizards, must negotiate complex environments.

## Background

The life and death of an organism often depends on its ability to perform well at some ecologically relevant task. Yet despite this significance we have little idea how well species are optimised for competing locomotor performances (1). Most scientists generally accept that the ability for natural systems to become optimised for a specific task is limited by structural, historic or functional constraints (performance trade-offs) (1-3). Climbing lizards provide a good example of constraint where the ability to climb requires the optimization of conflicting tasks such as speed, stability, or efficiency to maximize the animals’ fitness (3-7).

To explore this idea, we attempted to reconstruct the performance landscape of climbing locomotion using a 10 degree of freedom (DOF) robot based upon the lizard bauplan, including an actuated spine, shoulders, and feet, the latter which interlock with the surface via claws. This design allows us to independently vary speed, range of motion, foot angles, and movement kinematics, while simultaneously collecting data on climbed distance, stability and efficiency (Extended Data Movie S1). We can then compare the results from our robotic model to extant species to understand where constraints or trade-offs might occur, and if so, which performance characteristics different species may optimize for while climbing vertical surfaces.

We address three fundamental issues related to climbing locomotion: [1] How does speed, stability and efficiency interact to produce an optimal movement speed while climbing, [2] How does altering the foot angles of the fore and hind limbs, independently affect speed and stability, and [3] How does altering the range of motion of the spine and limbs independently, influence speed, stability and efficiency. This study will therefore demonstrate how robots can not only be inspired by biology, but how bio-inspired robots can contribute to our understanding of evolutionary change (8, 9).

## Methods

### Development of the bio-inspired climbing robot

We developed an agile climbing robot capable of climbing rough or compliant, vertical, and inclined surfaces based on the running trot of climbing geckos (Extended Data Figure S1). It is driven via lateral flexion, with diagonal pairs of limbs contacting the surface in unison following biomechanical patterns in lizards (10, 11). Each foot of the robot uses a 4-bar linkage system to simultaneously raise and push, or lower and pull the foot to engage the claws, mimicking the directional dependent adhesive systems of insects and lizards (12).

To minimise control complexity and ambiguity, we reduced the joints and their DOF in the robot to the minimum number for performing the climbing gait (Figure 1b). The robot has 10 servo motors: Two located along the trunk to allow spine flexion (2x Savöx SH-0257 mg, torque=2.2 kg.cm, USA; red joints in Figure 1b), four located in the shoulders to protract and retract the legs forward and backward (blue joints in Figure 1b), and four mounted in each leg to lift the feet (8x adafruit micro mg, torque=1.8 kg.cm, New York City, USA; yellow joints in Figure 1b). The mount design of the four foot servos permits the angle in which the feet are lifted to be rotated relative to the shoulder. This mimics the foot angle orientation in lizards (Figure 2f), which does not align with the body axis (13, 14). To increase stability on vertical surfaces, hip height is reduced among climbing lizard species compared with terrestrial species, to reduce centre of mass (COM) height (5). The electronic components were similarly arranged to centre and lower the COM, and to further reduce the overturning impulse moment, a simple abstracted rigid and static tail, 50 % the length of the robot, was added as a third contact point. Claw angles and leg lengths of the robot were optimized by iteratively testing variation in each and choosing dimensions which maximised the total distance moved forward.

**Figure 1:**
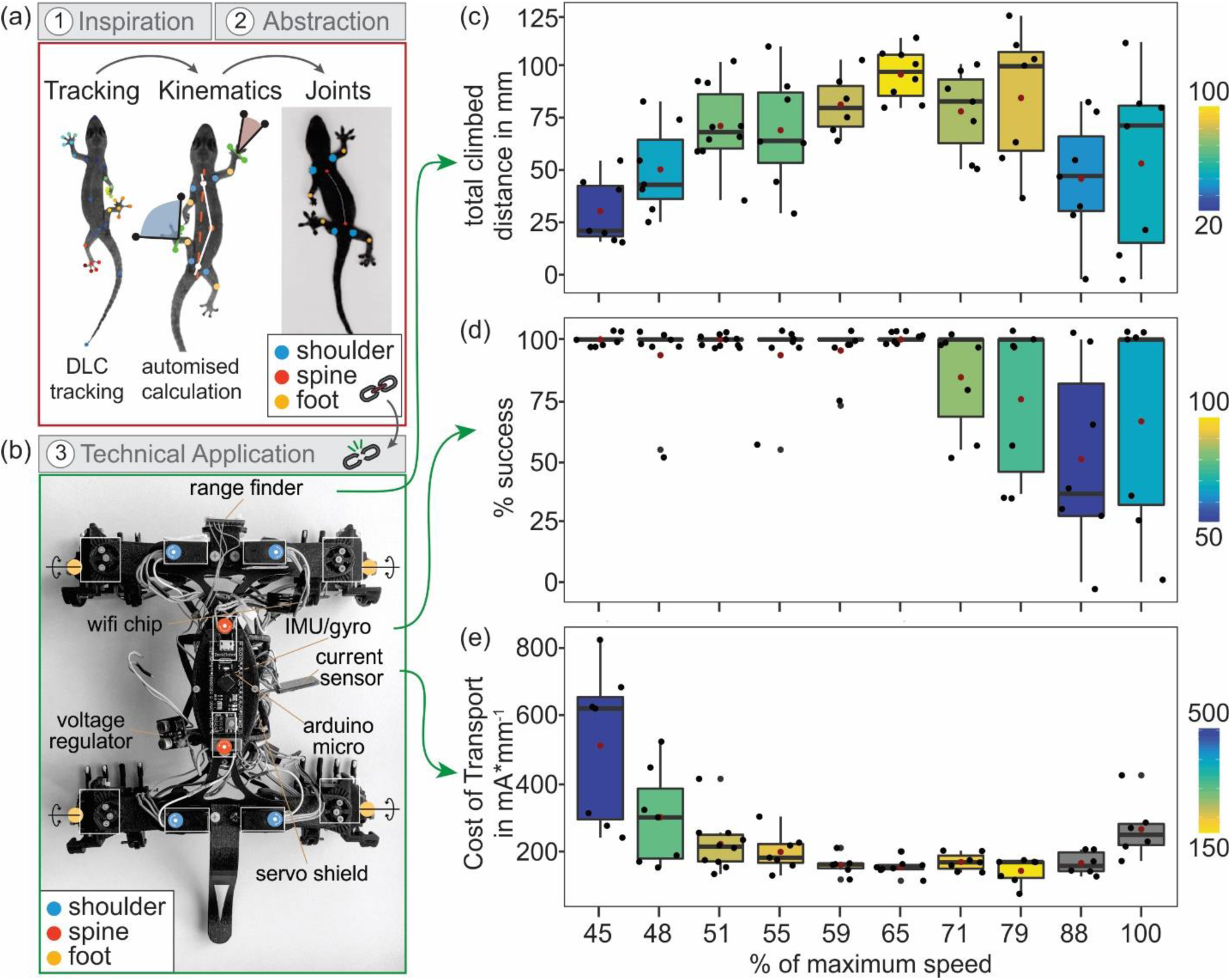
Speed, stability, and efficiency. (a) Illustrates the first steps of the biomimetic workflow: Inspiration and abstraction to design a bio-inspired robot by filming, tracking, and kinematic analysis. (b) Transfer of the derived design criteria. We developed a bio-inspired robot that mimics the lizards’ climbing locomotion and the basic morphology with 10 DOF together with sensors to measure climbed distance, stability of climbing, and the efficiency of the motors. (c-e) Results for the total climbed distance, the percentage of successful strides (∼ stability), and the cost of transport (∼ efficiency) with changes in input speed (as % of robot maximum speed.

**Figure 2:**
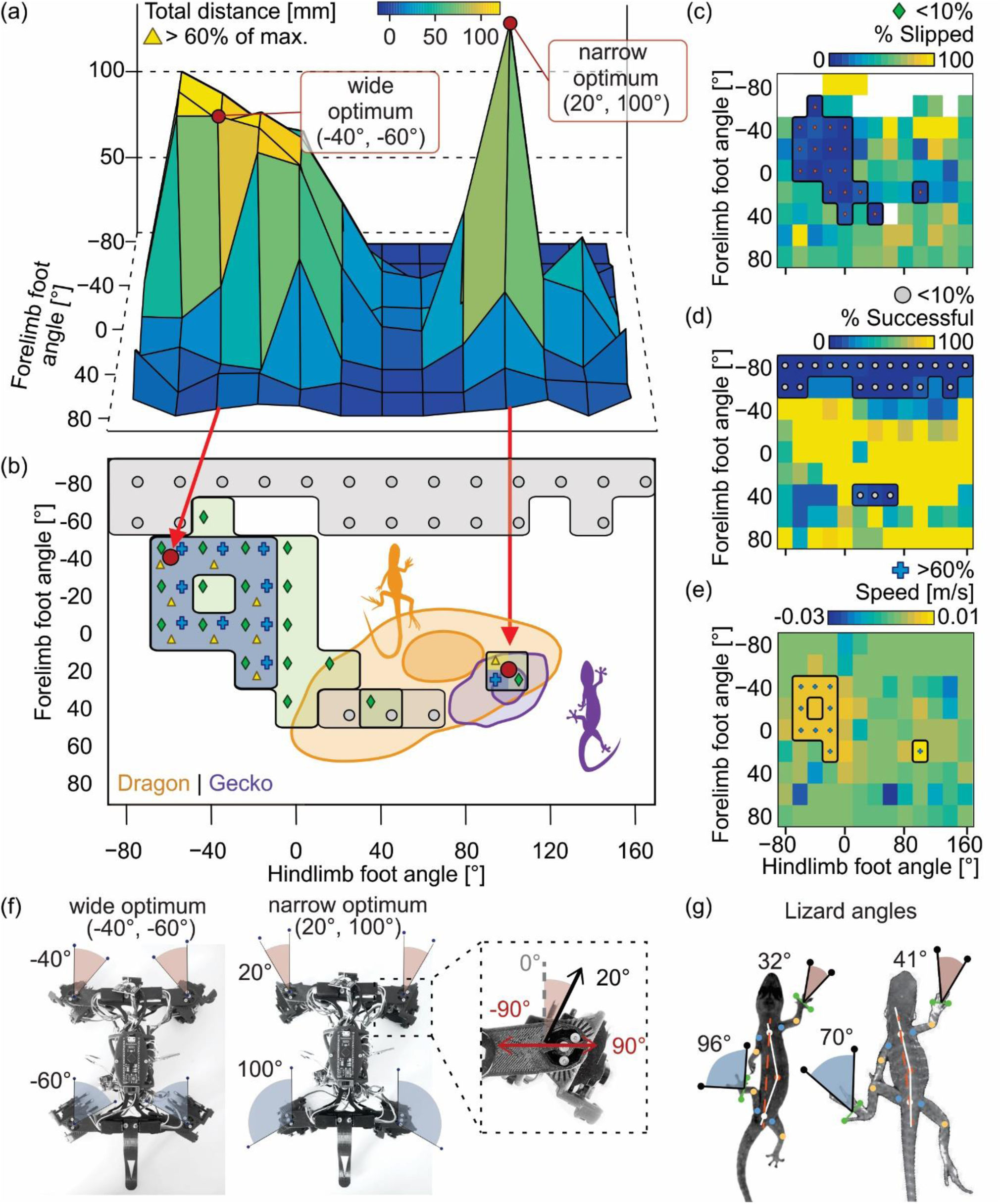
Foot angle performance landscape. (a, c, d, e) Show the resulting performance landscape of Total climbed distance (in mm), Success Percentage, Slipping Percentage, and speed (m/s), for the robot climbing with different fore and hind foot configurations (n = 444). (b) Foot angles during midstance for water dragons (*Intellagama lesueurii*; Orange; n = 127) and geckos (*Hemidactylus frenatus*; Purple; n = 384), with contours representing density estimates containing 70% and 20% of strides. The geckos overlay the narrow robotic optimum (20° fore, 100° hind). The climbing configuration for both the narrow and wide optimum are illustrated in (f), and mean angles for geckos and dragons are shown in (g).

For statistical accuracy and repeatability, we built and tested three third generation robots (X3) and one generation four robot (X4). The X3 generation robot weighed 332.7 g ± 3.2 g, and measured 240 × 180 × 65 mm (LxWxH). The X4 generation robot had minor hardware changes, with modifications to the bearing used along the spine joint, to decrease off-axis movement, and an additional sensor (see below). It weighed 330.6 g and measured 280 × 180 × 65 mm.

#### Electronics

The robots were built using 17 (X3) or 18 (X4) electronic components, which can be broadly categorized into power supply, motors, sensors, MCU and communication. The robot was powered by an onboard 7.4 V (2S) LiPo turnigy nano-tech battery (Hobbyking, Australia). Voltage from the battery was regulated to 5.5V by a 3A Step-Down Voltage Regulator (Texas Instruments, Lm2596, USA) in the X3 model or a Polulu 5V, 5A Step-Down Voltage Regulator in the X4 model (Polulu, D24V50F5, USA). The main processing unit was an Arduino micro (Deek robot, Atmega168P-AU, Shenzhen) which ran control software to coordinate the movement of servo motors, collect sensor data and to send/receive data to an ESP8266 Wi-Fi Chip (Espressif Systems, Shanghai, China). The ESP8266 opened a soft Access Point (AP), where a user was able log in to call the robots web interface for controlling gait parameters and downloading data after trials. The 10 servo motors were connected to a PCA9685 servo shield (Adafruit, New York), which provides power and control for all motors.

Two sensors were used in the X3 robot, three in X4, to collect data on distance climbed, orientation and current draw. Climbed distance during each stride was measured using a VL53L0X laser distance sensor (STMicroelectronics, Switzerland) mounted at the front of the robot. To measure orientation in space a MPU5060 (InvenSense, USA) Gyroscope was placed ventrally, along the body segment of the robot. The X4 generation robot was also able to measure the current draw of the motors using an INA219 current sensor (26 V ± 3.2 A max, adafruit, New York City, USA), mounted to the body segment. All sensors communicated with the Arduino via I2C.

#### Programming

The code for the robot was written in C++ and the Arduino communicated with the ESP8266 Wi-Fi Chip via UART-serial communication. After initial setup to calibrate sensors and move the robot into a default home position (Figure 1b), the robot waits until it receives a set of kinematic parameters, sent via the web interface. Each climbing step was carried out in a for loop, which was divided in 50 increments. In each increment eight motors were moved simultaneously; two motors lifted diagonally opposite feet off the wall while the remaining six performed limb and spine movement. The motors were controlled by a *moveMotor(X, A)* function, which moved a motor X to a desired angle A:

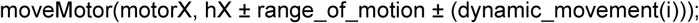

The desired angle A was not directly prescribed, but could be user prescribed: First the previous angle of the motor was calculated, onto which then the new increment is added: *Hx* set the middle angle of the limb motors by default to 90 degrees and aligned the spine motors. Afterwards the deflection of the home angle from the end of the previous step was added or subtracted by *range_of_motion*, the sign determining the direction of deflection. Out of this position the *dynamic_movment (increment i)* function was called to calculate the new angle to add to the previous. The dynamic_movement function determined the course of the motor angles to follow a sigmoidal curve during a step (50 increments). Every increment ended by a fixed delay given in milliseconds. This delay (*speed_val* variable) changed the speed of a robot’s movements (Extended Data Figure S1).

The robot was programmed to perform 11 strides in a row, with the first stride consisting of a half stride to move the robot from the default home position (limb motors at 90 degrees and both spine motors aligned), to the start of the stride pattern. The robot continued to climb until either the 11 steps were executed, it was <= 85 mm from the ceiling, or it fell off the wall.

#### Modulation

All parameters which can be modified via the web interface are listed and described in Extended Data Table S1. The following parameters were adjusted for this study;

##### Driving speed

(“Speed”) of the robot was modified by modifying the delay in ms added to the motor movement between every increment to stall before reaching the target angle. Delay was varied between 0 ms (maximum speed of the robot) to 18 ms in steps of 2 ms.

##### Foot angles

The foot angles were changed via hardware modification by adjusting two screws located within the fore and hindlimb foot joints (Fig. 2f, Fig. S3b, MovieS2). This joint is the attachment point for the servos which function to raise and lower the 4-bar linkage which in turn acts to interlock with the surface. Therefore modification of this angle results in the direction dependent claws acting in a different orientation relative to vertical.

##### Limb and Spine Range of Motion (ROM)

The input ROM set via the web interface defines the angle by which motors will be deflected in each direction from the default home position, and hence equals half of the resulting ROM in the gait (Movie S3). The design of the robot does not allow input values greater than 45°, as limbs would collide during a stride. Limb ROM angles are passed to the shoulder servo motors (blue joints, Figure 1b), while spine ROM angles are passed to the mid body servos (red joints, Figure 1b). The flow chart of the code for the robot is displayed in Extended Data Figure S5.

### Experiments and data analysis

The climbing robot was tested and filmed on a carpeted surface of 90° incline (i.e. vertical orientation), with a white corflute™ top as a reference point for distance measurements (Movie S4). For each configuration, the robot was given three attempts to climb the surface, with the results of the attempt downloaded following each trial. After every third run the battery of the robot was exchanged and recharged. Apart from the two modified parameters per trial set, the robot was tested in a default configuration (Table S2). For every trial set, all runs were then summarized by combining the 11 strides of each run to calculate distance climbed, stability and efficiency.

Two estimates for stability were included, the number of successful steps (%success) and the number of slipped steps (%slipped). The success rate (%success) was calculated as the fraction of strides the robot remained attached to the wall.

Unsuccessful strides (when the robot fell off the wall) were indicated by distance values higher than 80 cm or by sudden direction changes in the pitch detected by the gyro. The slip rate (%slip) was calculated using negative distance scores, based on the reasoning that if the distance to the target was greater than the previous step, the robot slipped. The %slip value therefore describes the percentage of negative steps within a trial.

Successful trials were analysed further to calculate distance and efficiency. Distance estimates were compiled using the mean distance between strides (including negative slip trials), as well as the total forward distance, a sum of all positive distance strides. Mean resultant speed was calculated using the mean distance between strides, divided by the time difference between strides. The cost of transport (COT) was calculated by dividing the mean current draw [mA] by the mean climbed distance [mm]. For graphical representations of COT only positive values were included, and values were log transformed to approach a Gaussion distribution.

#### Animal trials

384 steps resulting from 65 videos of 6 individual Asian House Geckos (*Hemidactylus frenatus*; mass = 3.19g ± 0.57g, SVL = 43.0mm ± 3.1mm), as well as 127 steps from 34 videos from four individual Water Dragons (*Intellagama lesueurii;* mass = 5.65g ± 0.99g, SVL = 53.8mm ± 3.23mm) were included. All lizards were wild caught and transported in cotton bags to the Animal Ecology Laboratory at the University of the Sunshine Coast under animal ethics permit ANA16104. Lizards were kept in glass terrariums and supplied with a daily refill of water and crickets as food. A natural day-night-cycle was maintained, the temperature was kept at 25° ± 5° and a relative humidity of 47.69 % ± 2.78 %.

During trials geckos were filmed climbing a white corflute (smooth plastic) vertical racetrack, while marine carpet was used for dragons. Videos were recorded with a high-speed camera (Fastec IL3 + Nikon 28 mm lens, 1280 × 1024 px, 250 fps, 1/500s shutter speed). The racetrack was illuminated using bright LED-spot lamps. Only runs where body deflection (of body axis) was < 20 degrees from vertical head-up were included in the analysis. The body axis was defined as the line passing through *Shoulder* and *Hip* tracking points.

#### Tracking

A Deep Residual Neural Network using the toolbox DeepLabCut (15) (DLC) was trained to track 34 tracking points per frame for all videos (Extended Data Figure S3a). The resulting accuracy after 77,000 iterations of training after network refinement was 3.04 px error (geckos) and 3.98 px (dragons), which is close to human accuracy of 2.7 px error (16).

#### Kinematic Analysis

All calculations described required a likelihood of > 90 % of all labels needed for the calculation, before inclusion in the analysis. The angles calculated and the orientation of the coordinate system are displayed in Extended Data Figure S3a. Foot angles were calculated for mid stance and for each foot. Footfall and end stance phases were automatically determined, using a greater than or less than 5 px change in foot marker position as a threshold. This threshold value was tuned until plotted foot fall pattern diagrams looked reasonable with a duty factor of ∼0.5 (17, 18). The foot vector was defined as the vector perpendicular to a vector which spanned the outermost toes of each foot. The angle between this foot vector and the body axis was used to estimate the foot angle for each frame. The mean of the middle index of the stance phase ± 1 frame was used as a final estimate of foot angle, meaning only stance phases longer than 3 frames were included.

The limb vector was estimated using the Shoulder and Knee markers for every limb. The angle between this limb vector and the Shoulder-Spine or Hip-Spine vector, for fore and hind limb respectively, was calculated for each frame during the stride phase of the foot. The range of motion was then determined by subtracting the limb angle of the start of the stride phase from the limb angle at the end of the stride phase.

The spine ROM was estimated as the mean of the Spine-Shoulder ROM and the Spine-Hip ROM. For the former the *Shoulder* and *Spine marker* are used to create a vector, for the latter *Hip* and *Spine* markers are used. The angle between each vector and a vector perpendicular to the body axis is calculated per frame throughout stride phase of each foot. The angle at the end of the stride is subtracted from the angle of the beginning of the stride.

## Results and Discussion

Even though higher speed may be linked with greater performance, most species use speeds between 60-80 % of their maximum capabilities, a result of the speed-stability trade-off (6). This is especially true for animals moving on challenging substrates, like narrow beams and branches or when climbing vertical surfaces. At high speeds slipping becomes more likely, animals may therefore choose slower speeds to avoid mistakes, indicating a change in the selective fitness in certain performance parameters (19, 20). Alternatively, animals may also choose speed during foraging to balance the cost of transport (COT) against the energetic reward of prey capture (21, 22), yet it’s unclear how each of these factors may independently influence speed choice.

To explore this, we systematically varied the input speed of our robot from 100 % of maximal speed (determined by the motors capabilities) to 40 % of maximum speed, and measured climbed distance, stability, and efficiency. We show that much like the lizard data, high speeds above 70 % of max, result in reductions in stability (Figure 1d, Extended Data Figure S2). Further we can show that low speeds result in an increase in the COT (Figure 1e). As a result of slipping, our robot was able to travel the furthest distance at intermediate input speeds (Figure 1c). This demonstrates the trade-off between these competing performance characteristics in a single system.

Speed, and stability might also be expected to vary with adhesion strategies used in climbing animals. Direction dependent adhesives and claws adhere to vertical surfaces by interlocking when pulled in one direction and detach easily when pushed in the opposite direction (12, 17). Since gravity acts in a consistent direction, the angle at which a lizard is climbing, and the angle of the feet relative to the body, will determine whether each foot is in a pulling or pushing configuration. For example, during head up climbing, if all feet are parallel to the body, then both the fore and hind limbs should be in the pulling configuration, providing passive stability. Yet geckos, other climbing lizards and many insects (e.g. cockroaches) show foot angles non-parallel to the body, and lateral forces can equal up to 50 % of the vertical forces (13, 14). With our robot we were able to independently modify both fore and hind limb foot angles to produce performance landscapes to show which configurations maximise speed and stability (Figure 2, Extended Data S3, Extended Data Movie S2). The foot angles of geckos and dragons were tracked and plotted onto the robot’s results.

Two distinct local optima for total forward climbed distance were observed (Figure 2a). The first was a broader, but lower optimum where both fore and hind foot angles show radial deviation (towards the body) of between −40° to 0° fore and between −60° to − 40° for the hind feet, with a mean climbed distance of 121 ± 10 mm. Yet a second, narrower, but higher optimum was also found with a foot angle configuration, showing ulnar deviation of 20° fore, and 100° hind resulting in a higher climbed distance of 186 ± 14 mm (Figure 2f). Both optima for distance were also stable. In both cases the percentage of successful climbs was at 100 % (Figure 2c, d), and both configurations exhibited minima for slipping (strides resulting in negative distance), suggesting no trade-off between both (Figure 2c). Of the two configurations, the wider optimum had slipping percentage of 5 ± 1.7 %, while the slipping percentage for the narrow optimum was 6.6 ± 6.6 %.

The results for the Gecko overlay exactly with the narrower, more confined robotic maximum (Figure 2b). The geckos’ mean forelimb showed ulnar deviation of 32° (95 % CI’S 30.6° – 33.5°), and the hindlimb 96° (95.1° – 97.3°; Figure 2g). Results for dragon forelimb showed ulnar deviation of 41° (34.7° – 46.9°), and the hindlimb 70° (64.8° – 74.6°), suggesting this largely terrestrial species occupies a similar but less optimal region in the climbing landscape. Both the narrowness of the robotic peak, and the low variation in angles among geckos suggest that there is a significant cost to deviating from this configuration for climbing species. We show that the forelimbs are mainly used in a gripping configuration, but this is less important for the hindlimbs, which instead appear to be more important to stabilize lateral undulations (13, 14). The reduction of safety and speed for the robotic model in the purely adhesive configuration (e.g. 0,0 degrees with both fore and hindlimbs parallel to the body) highlight the importance of lateral forces to maintain stability on inclined surfaces, and demonstrates how these may eliminate the trade-off between speed and stability (23).

Both species appear to avoid the wider optimum which provides at least similar levels of safety and speed. This may indicate a potential morphological constraint, where the structure and arrangement of muscles, tendons and bones in the fore and hind limb prevent adequate rotation of the foot. This might prevent the entire trait space from being explored among climbing species, similar to morphological constraints which have been demonstrated in other systems (24). Alternatively, it may be liked with habitat shape. The wider optimum may only be present on flat, or concave surfaces and is unlikely to be present on convex surfaces such as tree trunks or narrow perches like a branch. A convex surface would promote adhesive friction when limbs are rotated outwards following lateral undulation, while the opposite would be true on concave surfaces (25). In any case, without the performance landscape produced from the robotic model, the presence of an alternative performance peak, may have been difficult to observe.

Along with changes in the foot angle, changes in the limb and spine kinematics may be important for speed, stability, or efficiency. Speed in sprawling animals can be driven primarily by lateral bending of the trunk, or by movement of the limbs protracting and retracting with each step (26, 27). Considerable variation occurs among sprawling species as to the relative importance of each, with different lizards employing qualitatively different amounts of spine bending vs limb propulsion (27, 28). Multiple hypotheses have been developed to explain the role the spine might play in sprawled locomotion: it may speed up locomotion (4, 29, 30); increase turning ability (17, 31); stabilize the body to prevent slipping (13, 32); or alternatively enhance energy efficiency while moving (33). However, it is difficult to investigate this variation since changes in spine use is often coupled with variation in limb development making comparative studies difficult (11, 27).

Using our robot, we were able the vary the range of motion (ROM) of both the spine and limb independently and determine the influence on climbing ability for each combination. Further, given that these are likely to interact in a complex fashion with gripping ability, we tested locomotor ability of our robot on the ground as well as a vertical surface. The spine and limb ROM angles were also tracked and analysed for climbing geckos and dragons. We further plotted results for four species of walking squamates (34) onto the robotic landscapes.

On the ground the best distances are all reached in a diagonal band suggesting high speeds can be created with a number of combinations of the limb and spine (Figure 3a, Extended Data S4a). The fastest configuration (resultant speed = 0.112 m/s) appears in the lower left-hand corner in Extended Data Figure S4b, which minimises spine contribution, but maximises limb range of motion (Spine = 20°, Limb = −90°).

**Figure 3:**
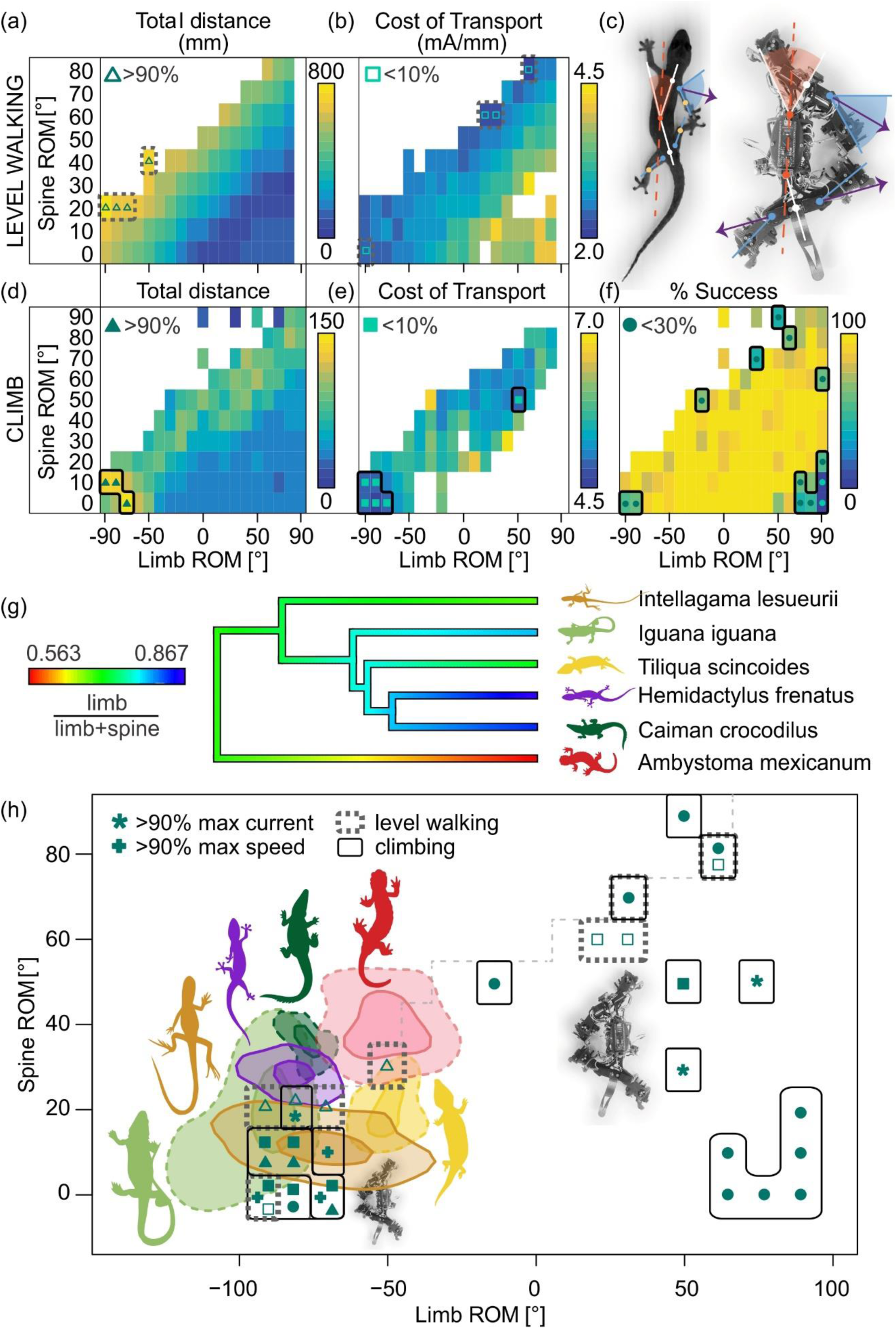
Spine and limb range of motion (ROM) performance landscape. Total forward distance (a) and cost of transport (b) for level walking (n = 238), for ROMs defined in (c). d-f show the total forward movement, cost of transport and %Success, for climbing trials respectively (n = 318). (g) Phylogenetic relationship of species included in (h), based upon (40). Colours represent the relative ROM of the limbs relative to the spine. (h) Comparison of the spine and limb ROM for both climbing geckos and dragons, alongside level walking for salamander, skink, iguana and Caiman reported by (34). Contours represent density estimates containing 70% and 20% of strides. Maximal regions for distance (Δ), speed (+) and current use (*) are indicated, along with minimum COT (□), and %success (o). Solid shapes, and solid outline represent climbing data, with open shapes representing level walking.

Here limb ROM is protracted forward during lateral flexion away from the limb, meaning both can contribute to increase the length of the stride (Extended Data Movie S3). Moving up and right along the diagonal, there is a slight decrease in speed, which flattens out once the limbs contribute less and the spine contributes more (e.g. resultant speed = 0.084 m/s, Limb at 0°, Spine 50°). Along this diagonal, additional increases in spine flexion are negated by an increasingly retracted limb at the beginning of the stance phase reducing any increase in stride length. This pattern is reflected in the robot’s current consumption with greater strides requiring greater currents, but when the COT data are explored the most economical gait appears to be a spine driven model (Spine = 80°, Limb = 60°), where limbs are highly retracted before foot down, since this likely reduces torsion between the foot and the surface (Figure 3b).

When climbing up the vertical substrate a similar pattern develops for distance and speed, with a diagonal of cases performing best, and maximal performance being located in the bottom left corner (Figure 3d, Spine 10°, Limbs −70°). However, in this regime, the robot was close to an unstable region, illustrating another potential trade-off between speed and stability (Figure 3f). This area also showed a relatively high current consumption (Extended Data Figure S4g), though correcting for distance covered using the COT, shows that both this corner, and a second optimum with moderate movement of the spine and limbs (∼Spine 50, Limb 40) both were areas of reasonable economy (Figure 3e).

The similarity in optimality maps between terrestrial and arboreal locomotion might suggest few major shifts in kinematics are required to move between these habitats, at least when considering the performance characteristics of speed, stability and efficiency. Such a result may explain why within species changes in kinematics were not observed in geckos (35) and why among species changes, may be relatively subtle (5). It may also explain why the transition between arboreal and terrestrial habitats occurs frequently in many different lizard groups (36).

Exploring the terrestrial and climbing data among sprawling tetrapods, there appears an evolutionary trend toward a limb driven model. Early tetrapod species, such as the salamander occupy a space using similar limb and spine contributions, which appears in a less favourable region on the robotic performance landscape (Figure 3h). Subsequent derived lineages of tetrapods appear to have moved down along the diagonal towards the robotic performance peak, reducing the movement of the spine, and increased limb ROM (Figure 3g).

The geckos clustered tightly in this lower left-hand corner, climbing using a limb driven gait (Spine 27.5°, Limbs −108°; Figure 3h). In some cases, geckos were able to outperform the robot, accessing unobtainable gaits by allowing the hindfeet to overstep the forefeet. Water dragons similarly occupied this left-hand corner zone, showing a limb driven model, though with a broader distribution of gaits, which overlapped those of the robot (Spine 14.7°, Limbs −96°).

All species appear to avoid the extreme lower left corner where stability start to decrease. The congruence between the evolutionary change in gait and the robotic performance landscape is compelling. While the extent to which the robotic performance landscape can replicate actual performance in biological systems remains to be resolved, the result does show the potential for robotic inspired biology to highlight potential selective forces driving the phenotypic change in organisms over evolutionary time.

Using our robotic model, we were able to create potential performance landscapes for a climbing lizard. This would not be possible using only animal data from extant species since movement kinematics in many lizards appear relatively invariant even with changes in climb angle (35), making exploring the parameter space difficult. Further computer simulations are limited because of the difficulty in accurately predicting model-environment interactions, like surface and material effects leading to a simulation-reality gap (37, 38). The extent to which these performance landscapes are transferable to animal models is unclear, but the convergence in optimal configurations suggest at least some broad conclusions can be drawn. Further, using this method we can show why certain phenotypes are not present among extant species, illustrating why these would be potentially mal-adaptive. These principles may be useful to compare with relative rates of evolution along differing evolutionary histories. It also highlights the importance of biological inspiration in application, particularly towards the optimization of industrial climbing robots, which like geckos, must negotiate complex environments (39).

## Acknowledgments

We thank David Labonte (Imperial College London), Robert Cieri (University of the Sunshine Coast), Taylor J. M. Dick (University of Queensland) and Michael Kasumovic (University of New South Wales) for helpful comments on draft manuscripts. This study was funded by an Australian Research Council Discovery Grant (DP180100220) awarded to CJC.

## Data and materials availability

Data and code used for all statistical analyses within this manuscript are available at https://doi.org/10.6084/m9.figshare.12325835.v1.

Code for automation of lizard biomechanics is available via github https://github.com/JojoReikun/ClimbingLizardDLCAnalysis. An illustration of gecko tracking, foot modification and ROM modification is provided in Supplementary Videos.

