## Extended Data for "Using a biologically mimicking climbing robot to explore the performance landscape of climbing in lizards"

**This SI file includes:**

Figures S1 to S5

Tables S1 to S3

Captions for Movies S1 to S4

**Other Supplementary Materials for this manuscript include the following:**

Movies S1 to S4

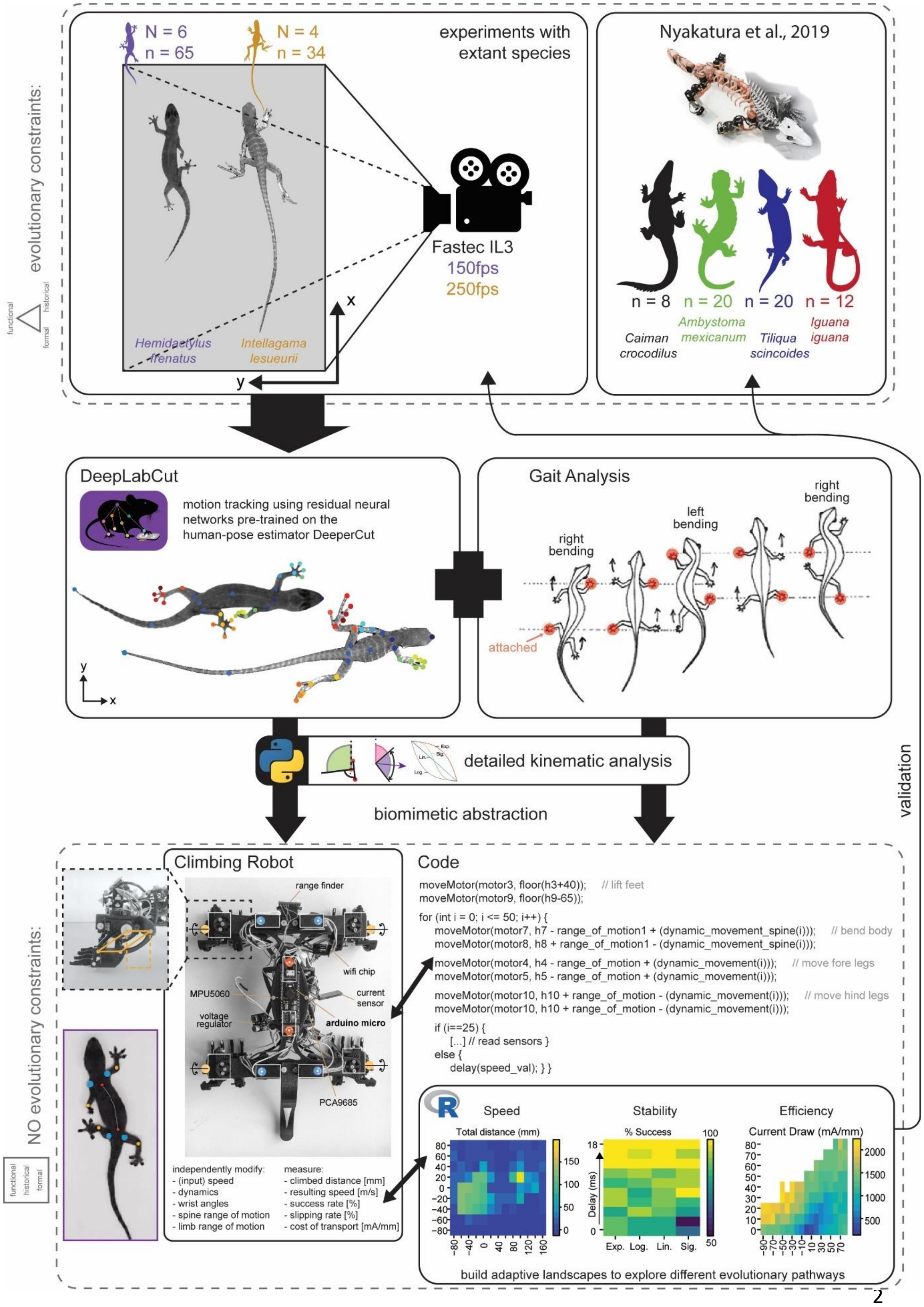

evolutionary constraints:
functional
historical
formal
N = 6
n = 65
N = 4
n = 34
experiments with extant species
Fastec IL3
150fps
250fps
Hemidactylus frenatus
Intellagama lesueurii
x
y
Nyakatura et al., 2019
n = 8
n = 20
n = 20
n = 12
Caiman crocodilus
Ambystoma mexicanum
Tiliqua scincoides
Iguana iguana
DeepLabCut
motion tracking using residual neural networks pre-trained on the human-pose estimator DeeperCut
Gait Analysis
right bending
left bending
right bending
attached
detailed kinematic analysis
biomimetic abstraction
validation
NO evolutionary constraints:
Climbing Robot
range finder
wifi chip
MPU5060
current sensor
voltage regulator
arduino micro
PCA9685
independently modify:
- (input) speed
- dynamics
- wrist angles
- spine range of motion
- limb range of motion
measure:
- climbed distance [mm]
- resulting speed [m/s]
- success rate [%]
- slipping rate [%]
- cost of transport [mA/mm]
Code
moveMotor(motor3, floor(h3+40)); // lift feet
moveMotor(motor9, floor(h9-65));
for (int i = 0; i <= 50; i++) {
moveMotor(motor7, h7 - range_of_motion1 + (dynamic_movement_spine(i))); // bend body
moveMotor(motor8, h8 + range_of_motion1 - (dynamic_movement_spine(i)));
moveMotor(motor4, h4 - range_of_motion + (dynamic_movement(i))); // move fore legs
moveMotor(motor5, h5 - range_of_motion + (dynamic_movement(i)));
moveMotor(motor10, h10 + range_of_motion - (dynamic_movement(i))); // move hind legs
moveMotor(motor10, h10 + range_of_motion - (dynamic_movement(i)));
if (i==25) {
[...] // read sensors }
else {
delay(speed_val); } }
Speed
Stability
Efficiency
Total distance (mm)
% Success
Current Draw (mA/mm)
Delay (ms)
Exp.
Log.
Lin.
Sig.
build adaptive landscapes to explore different evolutionary pathways


29

**Figure S1: Workflow for assessing lizards within the performance landscape.** Understanding performance constraints comparatively is difficult since it is unlikely that the entire performance landscape will be represented by extant species. Lizards were filmed and tracked using the DeepLabCut residual neural network, and movement kinematics were derived. A 10 DOF climbing robot was then constructed which was able to mimic the movement of these lizards, while being customisable to explore both kinematic and neuromechanical changes in gait. An array of sensors was able to collect data on competing performance criteria including speed, distance climbed, stability and efficiency, creating the performance landscape. Data from lizards, as well as published data on sprawling squamates was then overlayed onto this landscape to compare which performance criteria are optimised.

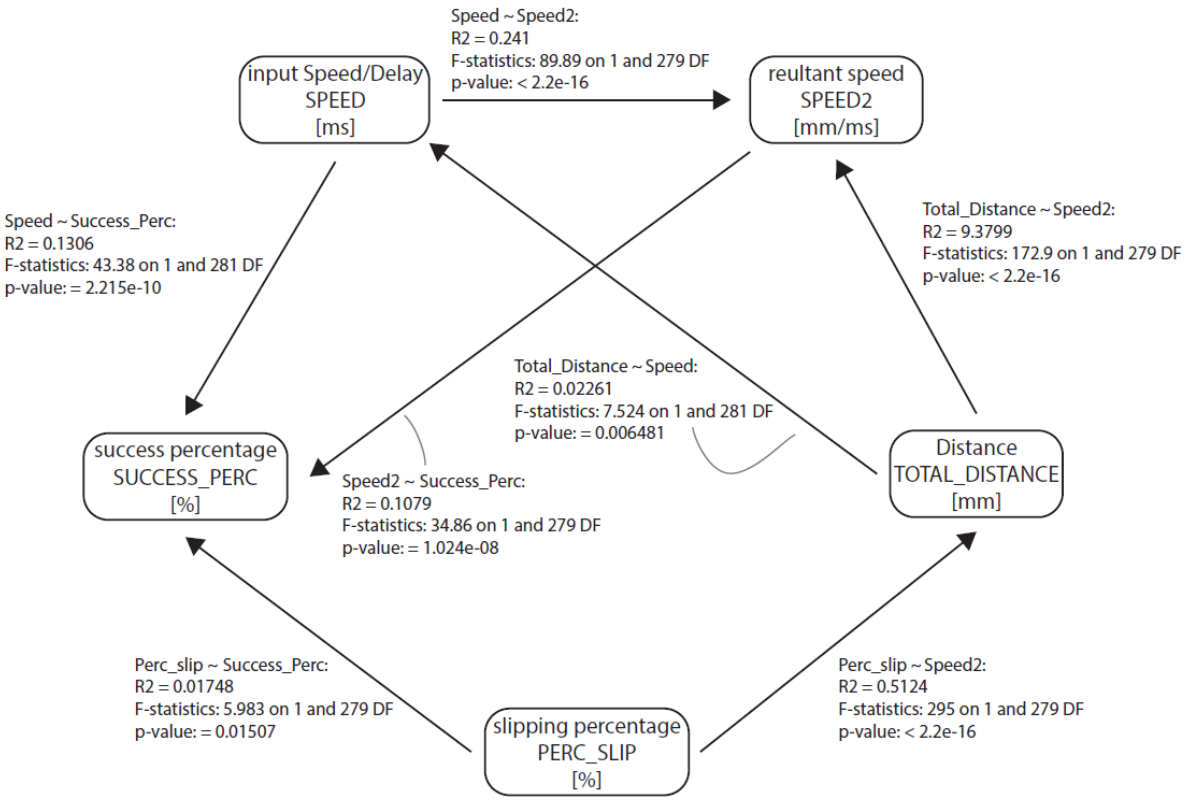


**Figure S2**: Statistical tests (anova) of influences between the variables: input speed, resultant speed, slipping percentage, success percentage, and total distance (parameter names as described in Table S3 in capitals).

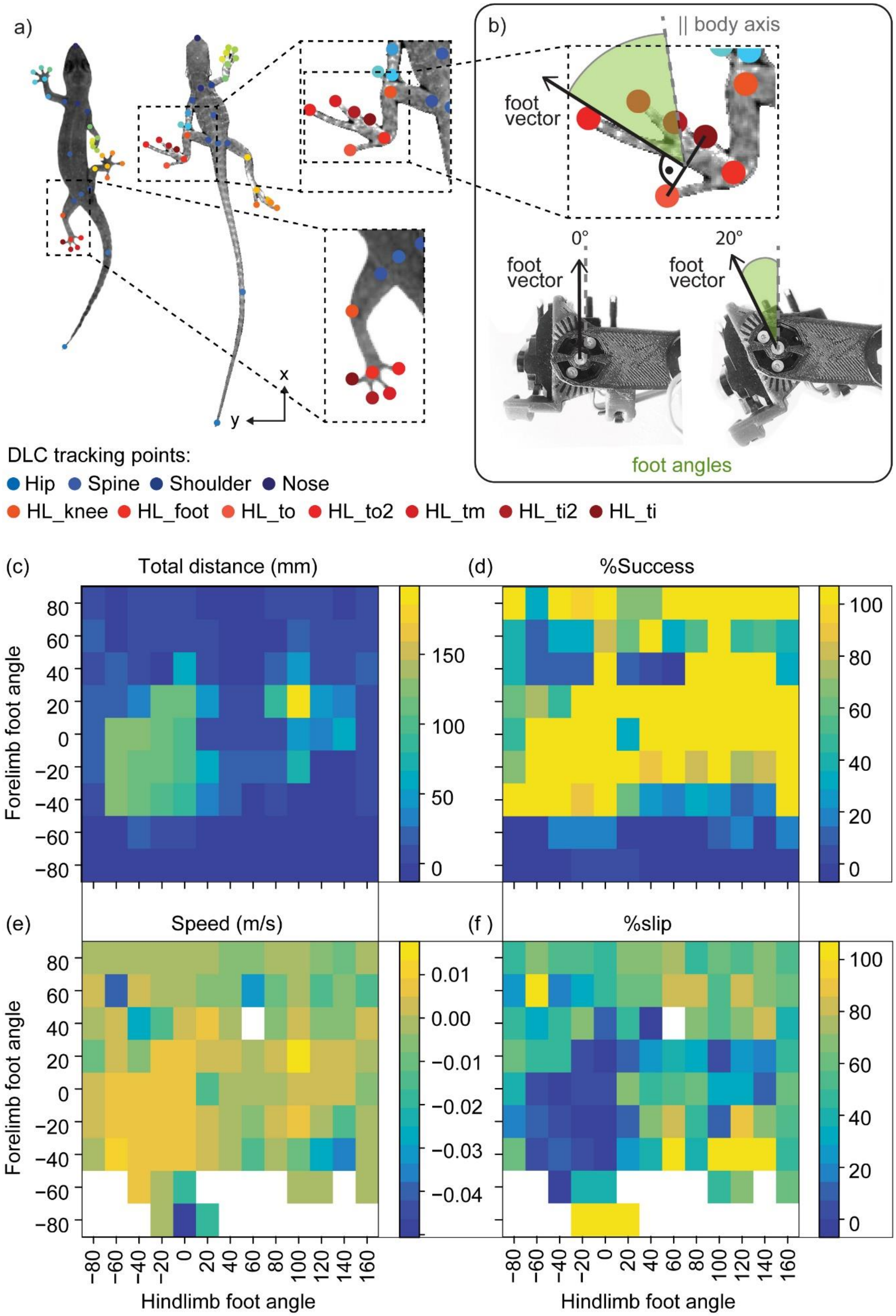
a)
b)
|| body axis
foot vector
foot vector
0°
foot vector
20°
foot angles
x
y
DLC tracking points:
Hip Spine Shoulder Nose
HL_knee HL_foot HL_to HL_to2 HL_tm HL_ti2 HL_ti
(c)
Total distance (mm)
(d)
%Success
(e)
Speed (m/s)
(f )
%slip
Forelimb foot angle
Hindlimb foot angle

**Figure S3: Detailed results of the foot angle performance landscape.** (a) Landmarks used in the calculation of fore and hind limb foot angles in lizards, along with (b) hardware modifications for foot angle changes in the robotic platform. (c-f) Performance landscape for forelimb and hindlimb foot angles in the robot, showing resulting Total distance climbed (mm), percentage of successful strides, resultant speed of robot (m/s) and the percentage of strides which slipped, respectively.

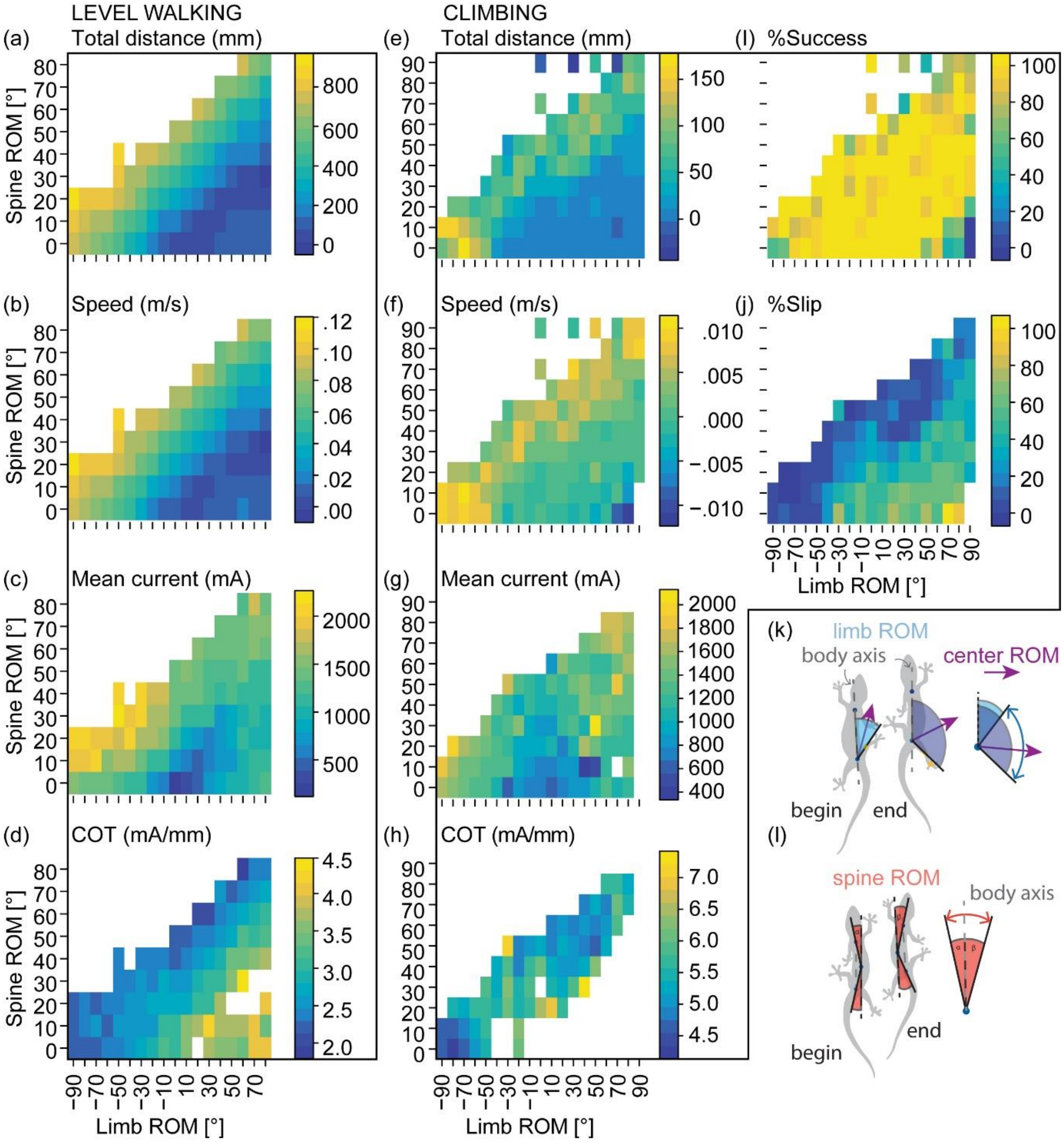


**Figure S4: Detailed results for the range of motion (ROM) performance landscape.** Heatmaps are shown for ROM values in Level walking (a-d) for total forward distance (mm), resultant speed (m/s), mean current consumption (mA) and cost of transport (mA/mm) respectively. Climbing performance results (e-j) for total forward distance (mm), resultant speed (m/s), mean current consumption (mA), cost of transport (mA/mm), percentage of successfully completed strides, and percentage of strides which slipped, respectively. (k-l) vectors used to calculate the ROM for the limb and spine respectively for lizards.

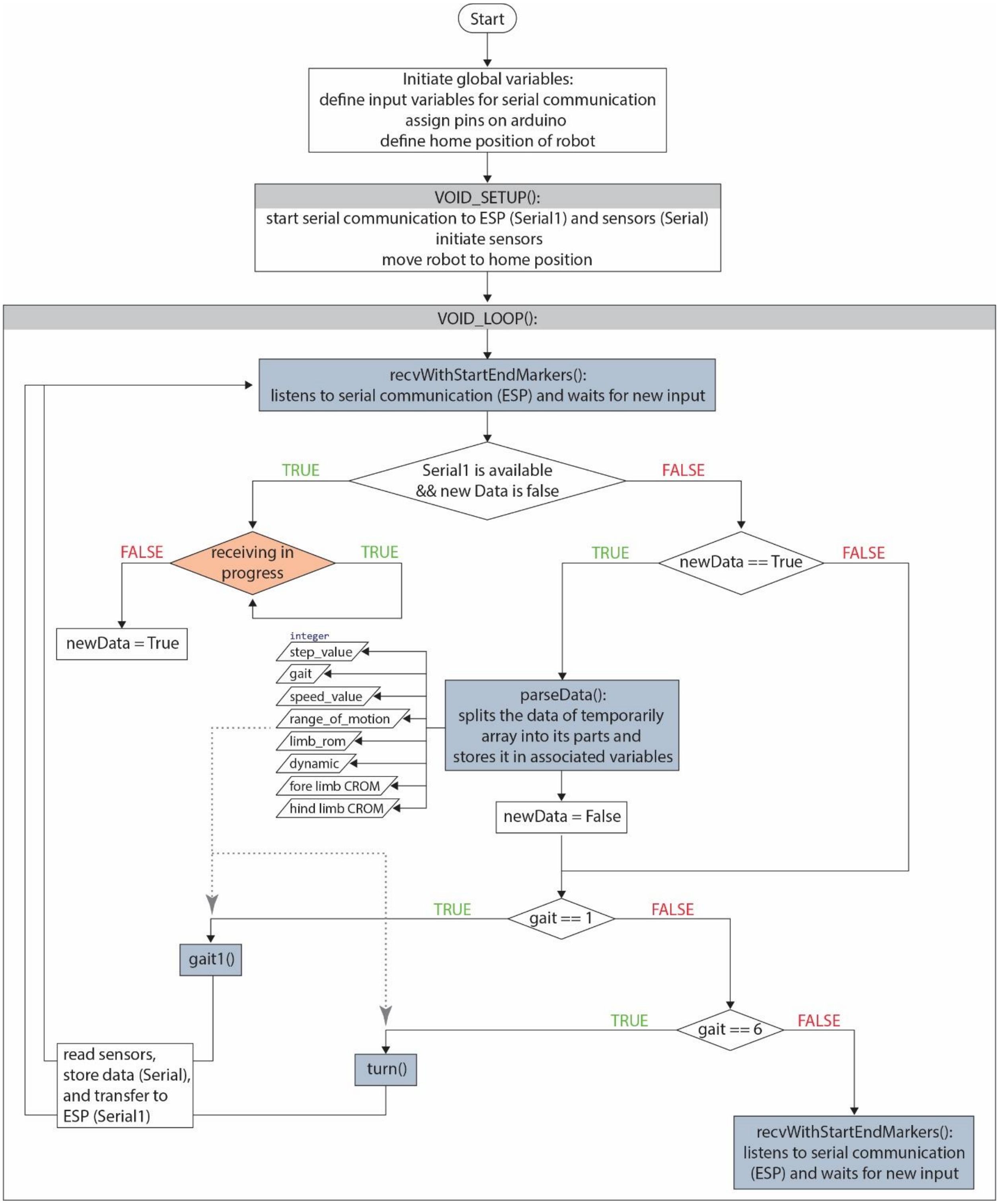

**Figure S5: Flowchart for the code which is used to control the climbing robot.** This code is uploaded to the Arduino microprocessor. After an initial set-up, the gait is configured and started via a website interface built up by the ESP WIFI chip. The gait, consisting of 11 steps, is executed, using the gait parameters defined by the user or using the default. Functions called are highlighted in blue. The orange diamond shape displays the communication with the ESP WIFI chip.

**Table S1:** Lists the software parameters that can be input via the robot's web interface to modulate the climbing locomotion and describes their function. Parameters that actively change the robot's gait are in white cells, parameters that are only stored in the csv file to simplify sorting trials are in grey cells.

| **Parameter** | **Function** |
|---|---|
| Step | The step to start the gait from. Default is 0 |
| Gait | Gait 1 = climbing gait. Further gaits are to be implemented, e.g. turning gait will be Gait 2 etc. |
| Speed | Defines the delay in ms added between each of the 50 increments of the motor movement to stall before reaching the target angle |
| Limb range of motion | Defines the target range of motion angle **r** for the limbs (blue joints, Fig 1b), by which the servo motors deflect in each direction. Resulting ROM is double the input value. |
| Spine range of motion | Defines the target range of motion angle **r** for the spine (red joints, , Fig 1b), by which the servo motors deflect in each direction. Resulting ROM is double the input value. |
| Robot | The name of the robot used for the trials. Each robot has its own website with its own default values flashed onto the ESP chip |
| Front Leg Angle | CROM for fore limbs |
| Hind Leg Angle | CROM for hind limbs |
| Fore foot angle | The fore foot angle as adjusted in the hardware of the robot |
| Hind foot angle | The hind foot angle as adjusted in the hardware of the robot |
| Toe Angle | The angle between the toes (spreading). Default is 0 |
| Claw Angle | The angle of attack of the claws compared to the surface. Default is 10 |
| Surface | The name of the surface used to cover the climbing test wall with. Default is "coarse carpet" |
| Angle of Inclination | The incline of the climbing test wall. Default is 90 degrees |

**Table S2:** Lists the conducted test series, the parameters changed for each, the robot(s) used for the trials and the number of trials.

| **Test series** | **Testing config** | **Changed parameters** | **robot** | **n** |
|---|---|---|---|---|
| Speed - stability - efficiency | Limb&Spine ROM = 25,<br><br>Foot angles = 0 (f), 0 (h) | Speed | X4_hulk | 284 |
| Foot angles | Limb&Spine ROM = 25,<br><br>Foot angles = 0 (f), 0 (h),<br><br>Speed = 5 | Fore-foot angle, hind-foot angle | X3_green | 444 |
| ROM climbing | Limb&Spine ROM = 25,<br><br>Foot angles = 0 (f), 0 (h),<br><br>Speed = 5 | Limb ROM, spine ROM (5 increments)<br>+ (1 increments) | X4_hulk | 238<br><br><br>+ 469 |
| ROM floor | Limb&Spine ROM = 25,<br><br>Foot angles = 0 (f), 0 (h),<br><br>Speed = 5 | Limb ROM, spine ROM | X4_hulk | 318 |

**Table S3**: Lists the parameters gained from the robot trials for each data set.

| **parameter** | **unit** | **description** |
|---|---|---|
| name | - | name of the csv file for the trial downloaded from the robot's website (follows the pattern: robot name (nr), e.g: X4_hulk (42)) |
| Total_Distance | mm | the total climbed distance calculated by taking the first value (first step) and subtracting the minimum value of all 11 steps |
| mean_Distance | mm | describes the mean distance of the individual distances climbed with each step |
| Mean_Time | ms | the time for each step is measured by the Arduino time library. Mean time is the mean for all steps |
| Success_perc | % | described the percentage of successful steps for a trial. Successful means the robot did not fall off the wall |
| strides | - | the number of steps achieved per trial |
| Temp | °C | mointors the temperature of the Arduino. Not used for analysis |
| Mean_Current_Consumption | mA | describes the mean current draw of the motors for a trial. The current draw is measured by the INA219 current sensor which is integrated in the robot's circuit |
| Current_Spike | mA | describes the maximum current draw value which occurred during a trial. Not used for analysis |
| Robot | - | the name of the robot. Following the pattern: Generation_name |
| Fore foot angle | ° | the foot angles of the fore foot compared to the body axis of the robot |
| Hind foot angle | ° | the foot angles of the hind foot compared to the body axis of the robot |
| Toe_Angle | ° | the angle between the toes (default 0°) |
| Claw_Angle | ° | the angle between the claw and the foot sole. This is also the perturbation angle of the claws to the surface |

| | | |
|---|---|---|
| Surface Incline | ° | inclination of the surface the robot trials were conducted on (default 90°) |
| Dynamic | - | the dynamic function, which the robot's servo motors follow. Four dynamics were programmed: Linear (4), Sigmoidal (3), Exponential (2), Logarithmic (1) |
| Dynamic3 | - | exchanged the numbers of the dynamic function with their actual name: e.g.: sigmoidal = sig |
| Speed | ms | the input speed equals the delay in ms which is integrated between the increments of motor movement. The motor movement to reach the defined goal angle is divided into 50. |
| Speed2 | mm/ms | the resultant speed equals the speed of the robot calculated by dividing the mean_distance by the mean_time |
| Limb_range_of_motion | ° | the input limb range of motion equals half the resulting limb range of motion. |
| Spine_range_of_motion | ° | the spine range of motion equals half the resulting spine range of motion. |
| Front_limb_angle | ° | the CROM angle of the fore limbs describes the angle of the fore limb compared to the perpendicular of the body axis from which the limb range of motion is executed |
| Hind_limb_angle | ° | the CROM angle of the hind limbs describes the angle of the hind limb compared to the perpendicular of the body axis from which the limb range of motion is executed |
| Mean_forward | mm | the mean forward climbed distance excludes all slipped steps in the mean distance |
| Perc_slip | % | the percentage of steps per trial with a negative climbed distance |
| COT | mA/mm | calculated by dividing the mean current draw by the total climbed distance divided by the number of achieved steps |

**Movie S1:** Bioinspiration process. Deeplabcut tracking results for dragons and geckos. The tracking accuracy (test error) of the networks was 3.04 px for the geckos and 3.98 px for the dragons.

**Movie S2:** Modification of the fore and hind foot angles independently to explore the performance landscape. Results are compared to the extant lizards, mean foot angles during midstance for are shown for water dragons (Intellagama lesueuerii; n = 127) and geckos (Hemidactylus frenatus; n = 384).

**Movie S3:** Climbing and walking with different limb and spine range of motion configurations. Configurations visualized are at the ideal ROM (-90°, 20°) with ROM overlaid. Also illustrates climbing under three different limb and spine ROM configurations; Only spine (0°, 50°), only limb (50°, 0°) and equal spine and limb (50°, 50°).

**Movie S4**: Various climbing configurations shown for robots of both the X3 and X4 generation. Also shown is the ability to climb on several inclines, including an overhang configuration.
